# Different isolates of *Leishmania aethiopica* elicit distinct effector functions in primary human monocytes

**DOI:** 10.1101/2025.02.26.640320

**Authors:** E. Cruz Cervera, E. Adem, E. Yizengaw, Y. Takele, I. Müller, J.A. Cotton, P. Kropf

## Abstract

In Ethiopia, cutaneous leishmaniasis (CL) is caused by *Leishmania* (*L*.) *aethiopica* and presents as a spectrum of clinical forms ranging from self-healing to persistent, disfiguring lesions.

Monocytes are some of the first cells to encounter *Leishmania* parasites in the host. Despite this, their role in *L. aethiopica* infection has not been investigated. In this study, primary human monocytes were co-incubated with different *L. aethiopica* isolates – three isolates recently collected from CL patients and one long-term cultured isolate – and their main effector functions were evaluated.

After 2 hours of co-incubation, over 30% of monocytes associated with all four *L. aethiopica* isolates, with significantly higher association with recently isolated *L. aethiopica* than with the long-term cultured parasites. Phagocytosis of *L. aethiopica* parasites by monocytes was confirmed by confocal microscopy.

Co-incubation of monocytes with all L. aethiopica isolates resulted in upregulation of reactive oxygen species in monocytes.

Following incubation of monocytes with all *L. aethiopica*, five chemokines – monocyte chemoattractant protein (MCP)-1, MCP-4, macrophage inflammatory protein (MIP)-1α, MIP-1β, and interleukin (IL-8) – and four cytokines – tumour necrosis factor (TNF)-α, IL-1β, IL-6, and IL-10 – were detected.

Our results show that the interaction of monocytes with the long-term cultured *L. aethiopica* differ as compared to those recently collected from CL patients. Furthermore, we define an *in vitro* model for the investigation of monocyte effector functions that may be useful to elucidate the role that parasites and their interactions with monocytes play in the different presentations of lesions caused by *L. aethiopica*.

## INTRODUCTION

The leishmaniases are a group of neglected tropical diseases caused by protozoan parasites of the genus *Leishmania*. These diseases present as a wide spectrum of clinical manifestations, ranging from self-healing skin lesions to potentially fatal visceral infections. They are primarily classified into two main forms: visceral leishmaniasis (VL) and cutaneous leishmaniasis (CL). VL is characterised by the dissemination of *Leishmania* parasites to internal organs such as the liver, spleen and bone marrow [1, 2]. CL includes a variety of presentations where the parasite primarily infects the skin and mucosal tissue. CL is endemic in 90 countries, with 205,986 new cases reported by the WHO in 2022 [3]. Eight countries – Afghanistan, Algeria, Brazil, Colombia, Iran, Iraq, Peru and Syria – accounted for 85% of the global cases, and children under 15 years old represented 37% of CL cases worldwide [3]. The burden of CL includes permanent scarring and potential disfigurement, resulting in important psychological effects and social stigmatisation [4–6].

In Ethiopia, CL present mainly as three clinical forms [7, 8]:

- Localised cutaneous leishmaniasis (LCL), which is the most common form of CL, presents as one or more ulcerative skin lesions that develop at or near the site of the sandfly bite [7]. While LCL is typically self-healing, treatment is sometimes provided for persistent lesions [7].
- Mucocutaneous leishmaniasis (MCL) is a severe and debilitating form of CL that predominantly affects the mucosal tissues of the nose, mouth, and pharynx [7].
- Diffuse cutaneous leishmaniasis (DCL) is a chronic form of CL characterised by the widespread dissemination of non-ulcerating lesions [7, 8]. Both DCL and MCL require treatment [9].

According to the most recent WHO data, Ethiopia reported 1505 CL cases in 2023 [10]; however, this is likely to be an underestimate of the actual burden due to underreporting, as a result of limited resources, remoteness of endemic areas, and the lack of a system to report cases [11, 12]. The Ethiopian Ministry of Health reported an estimate of 20,000 to 30,000 yearly CL cases in 2013 [7]. The number of people at risk of CL in Ethiopia was estimated at 29 million [13].

*Leishmania* parasites are transmitted during the blood meal of female sandflies [14]. *L. aethiopica* causes most CL cases in Ethiopia [8]. Three *Phlebotomus* (*P*.) species have been identified as vectors for *L. aethiopica*: *P. pedifer* and *P. longipes* are considered to be the main vectors, with *P. sergenti* only identified once as naturally infected with *L. aethiopica* [15–17].

Since sandflies feed by lacerating capillaries to create a blood pool, monocytes are likely to be among the first cells to come into contact with *Leishmania* parasites [18]. Monocytes are circulating immune cells that make up ∼10% of peripheral blood cells in humans [19]. In the response to infection, monocytes are rapidly recruited and can perform a range of antimicrobial functions, including phagocytosis, production of reactive oxygen species (ROS), and the release of a range of cytokines and chemokines [19]. Furthermore, monocytes are able to differentiate into dendritic cells and macrophages [20]. In humans, monocytes are divided into at least three subsets, based on the expression of CD14 and CD16 [21]. The subsets have distinct functions that have been characterised by both functional and ribonucleic acid sequencing (RNA-seq) studies: classical monocytes (CD14^++^ CD16^−^) mainly play a role in phagocytosis, adhesion and migration; intermediate monocytes predominantly carry out antigen presentation (CD14^++^ CD16^+^); while non-classical monocytes (CD14^+^ CD16^+^) are principally responsible for inflammatory cytokine production, and complement- and Fc receptor-mediated phagocytosis [22–24]. However, RNA-seq studies have proposed different ways of classifying monocytes, often into more subsets, noting the heterogeneity within each of the three conventional subsets [25, 26].

Human monocytes, isolated from peripheral blood, have been shown to become infected by several species of *Leishmania in vitro* [27] and produce reactive oxygen species (ROS). Human monocytes were shown to kill *L. donovani* promastigotes in a hydrogen peroxide-dependent manner [28]. Hoover *et al.* showed monocyte killing of *L. donovani* amastigotes when interferon (IFN)-γ was added to cultures, although the mechanism of killing was not determined [29]. Another study showed IFN-γ induced killing of *L. donovani* by human monocytes, which correlated with increased hydrogen peroxide production [30]. Ritter and Moll demonstrated that the chemokine monocyte chemoattractant protein (MCP)-1 acted synergistically with IFN-γ to induce human monocyte killing of *L. major*, in a process correlating with increased superoxide production [31]. A more recent study showed that classical, but not intermediate and non-classical, monocytes killed *L. braziliensis* amastigotes in a ROS-dependent manner [32].

Most of the work on monocyte effector functions in response to *Leishmania* parasites has been done with parasite isolates that had been maintained in culture for an extended period of time and/or passaged *in vivo* in animals, but rarely with parasites recently isolated from patients. The only study assessing the interaction between blood immune cells and *L. aethiopica* was done with a parasite that was kept in culture for decades. Here, the main aim was to investigate different effector functions of human monocytes to three recently isolated *L. aethiopica* and compare those to long-term cultured *L. aethiopica*.

## MATERIALS AND METHODS

### Leishmania aethiopica parasites

Four isolates of *L. aethiopica* were used in this study: one long-term cultured isolate, (MHOM/ET/72/L100) [33, 34] referred to as *L. aethiopica* lab, and three isolates recently obtained from LCL patients in Lay Gayint, Amhara region, north-west Ethiopia, as described in [35, 36], referred to as clinical isolates *L. aethiopica* 1, 2 and 3. These parasites were grown from skin scrapings and were frozen as soon as they reached stationary phase. They were sent to the UK, where large stocks of frozen parasites were prepared for further use. Once thawed, the parasites were used for a maximum of 3 weeks.

Parasites were grown in M199 medium supplemented with 10 μM hemin, 1x Eagle’s minimum essential medium (MEM) vitamin solution, 25 mM HEPES, 0.2 μM folic acid, 100 µM adenine, 10% heat-inactivated fetal bovine serum (HI FBS), 8 μM 6-biopterin (Sigma-Aldrich, USA), 100 units/mL penicillin and 100 μg/mL streptomycin (Thermo Fisher Scientific, USA) and were incubated at 26°C. Stationary phase *L. aethiopica* were enriched for metacyclic parasites by eliminating non-metacyclic parasites by peanut agglutination, as described in [37]. Of note, the percentage of metacyclic parasites was significantly lower with *L. aethiopica* lab than each of the three clinical isolates (Table S1).

Parasites were stained using 1μM CellTrace™ Far Red dye (Thermo Fisher Scientific, USA). Over 98% of parasites were stained with the Far Red dye.

### Monocytes

Primary human monocytes were purified from the peripheral blood of healthy volunteers. 100-120 mL of blood was collected in heparin tubes (BD, USA) and monocytes were isolated by double gradient centrifugation [38]. The blood was first overlayed onto an equal volume of Histopaque®-1077 Hybri-Max™ (Sigma-Aldrich, USA) to collect the peripheral blood mononuclear cells (PBMCs), that were then overlayed onto an equal volume of 46% Percoll® solution. Monocytes were obtained from the interphase ring. All steps were performed in polypropylene tubes to avoid monocyte adhesion. Monocyte purity following purification was >80%.

2×10^5^ monocytes were incubated alone or in the presence of 2×10^6^ *L. aethiopica* parasites stained with Far Red dye (*L. aethiopica*^FR^). Incubations were performed in RPMI medium (Thermo Fisher Scientific, USA) supplemented with 5% HI FBS (Sigma-Aldrich, USA), 100 units/mL penicillin and 100 μg/mL streptomycin (Thermo Fisher Scientific, USA) in 5 mL round-bottom polypropylene tubes at 37°C, 5% CO_2_, for 2 hours. The supernatants were collected and frozen at −20°C until further use.

### Flow cytometry: monocyte association with *L. aethiopica* and ROS production

After 1 hour 45 minutes of incubation, cells were stained with anti-CD14^PE^ (clone M5E2; BioLegend, USA) and anti-CD16^eFluor™^ ^450^ (clone CB16; Thermo Fisher Scientific, USA) for 15 minutes. The different monocytes subsets were defined as follows: classical monocytes as CD14^++^ CD16^−^, intermediate monocytes as CD14^++^ CD16^+^ and non-classical monocytes as CD14^+^ CD16^+^ (Figure S1). Cells were then washed twice in PBS and were resuspended in 500 µL of ROS detection solution from the ROS-ID^TM^ Total ROS detection kit (Enzo Life Sciences, USA). Cells were incubated in ROS detection solution for a further 30 minutes at 37°C with 5% CO_2_, after which tubes were placed on ice and immediately processed. The percentage of monocyte subsets associated with each *L. aethiopica*^FR^ isolate and the ROS production were assessed by flow cytometry.

The percentages of live monocytes after the 2-hour incubation were measured by exclusion of Zombie Violet (BioLegend, USA) and were >98%.

Flow cytometry acquisition was performed using an LSR II (BD, USA) and data were analysed on FlowJo™ v10.10 Software (BD, USA).

### Confocal microscopy

3×10^5^ purified monocytes were incubated alone or in the presence of 3×10^6^ *L. aethiopica*^FR^ parasites for 2 hours, before being washed in PBS and transferred onto 24 mm x 24 mm glass coverslips (Thermo Fisher Scientific, USA) precoated with an excess of 0.01% poly-L-lysine. After 30 minutes at room temperature, the coverslips were washed with PBS and fixed with 2% paraformaldehyde (Sigma-Aldrich, USA). Twenty minutes later, the coverslips were washed twice in PBS and incubated with a rabbit anti-human CD14 IgG polyclonal antibody (PA5-13305, Thermo Fisher Scientific, USA) and incubated overnight at 4°C. The coverslips were washed with PBS and Alexa Fluor™ 488 conjugated goat anti-rabbit IgG antibody (A27034, Thermo Fisher Scientific, USA) was added and incubated for 1 hour at room temperature. Coverslips were then transferred to a 6-well plate and washed with PBS.

The coverslips were then placed over 10 µL of VECTASHIELD mounting medium with DAPI (Vector Laboratories, USA) on a glass microscope slide.

Slides were visualised under an SP8 LIGHTNIGHT confocal microscope (Leica Microsystems, Germany) with a 100x objective and images were acquired using the LAS X software (Leica Microsystems, Germany). 3-colour imaging was performed to assess staining with DAPI, CD14 staining of monocytes and the Far Red label of *L. aethiopica*. Images were acquired as z-stacks of at least 20 images to enable 3-dimensional visualisation. Images were analysed using Fiji software [39].

### Chemokine and cytokine measurements

The levels of the following chemokines: IL-8, eotaxin, eotaxin-3, interferon-γ-induced protein (IP)-10, monocyte chemoattractant protein (MCP)-1, MCP-4, macrophage-derived chemokines (MDC), macrophage inflammatory protein (MIP)-1α, MIP-1β and thymus- and activation regulated chemokine (TARC) and cytokines: IFN-γ, IL-1β, IL-2, IL-4, IL-6, IL-10, IL-12p70, IL-13, TNF-α were measured by multiplex assay using V-PLEX Proinflammatory Panel 1 and Chemokine Panel 1 Kits (Meso Scale Diagnostics, Rockville, USA).

### Statistics

Data were evaluated for statistical differences using Mann-Whitney and Kruskal-Wallis tests (GraphPad Prism 10). Differences were considered statistically significant at *p*<0.05. Unless otherwise specified, results are expressed as mean ± standard deviation (SD).

## RESULTS

### Association of monocytes with *L. aethiopica*

The ability of the different monocyte subsets to associate with *L. aethiopica* was determined. Results presented in Figure 1A-D and Table S2 show that all three subsets had the ability to associate with monocytes, with the classical monocytes displaying the highest mean association. Of note, not all monocytes associated with *L. aethiopica* (hereafter referred to as unassociated monocytes).

**Figure 1-.**
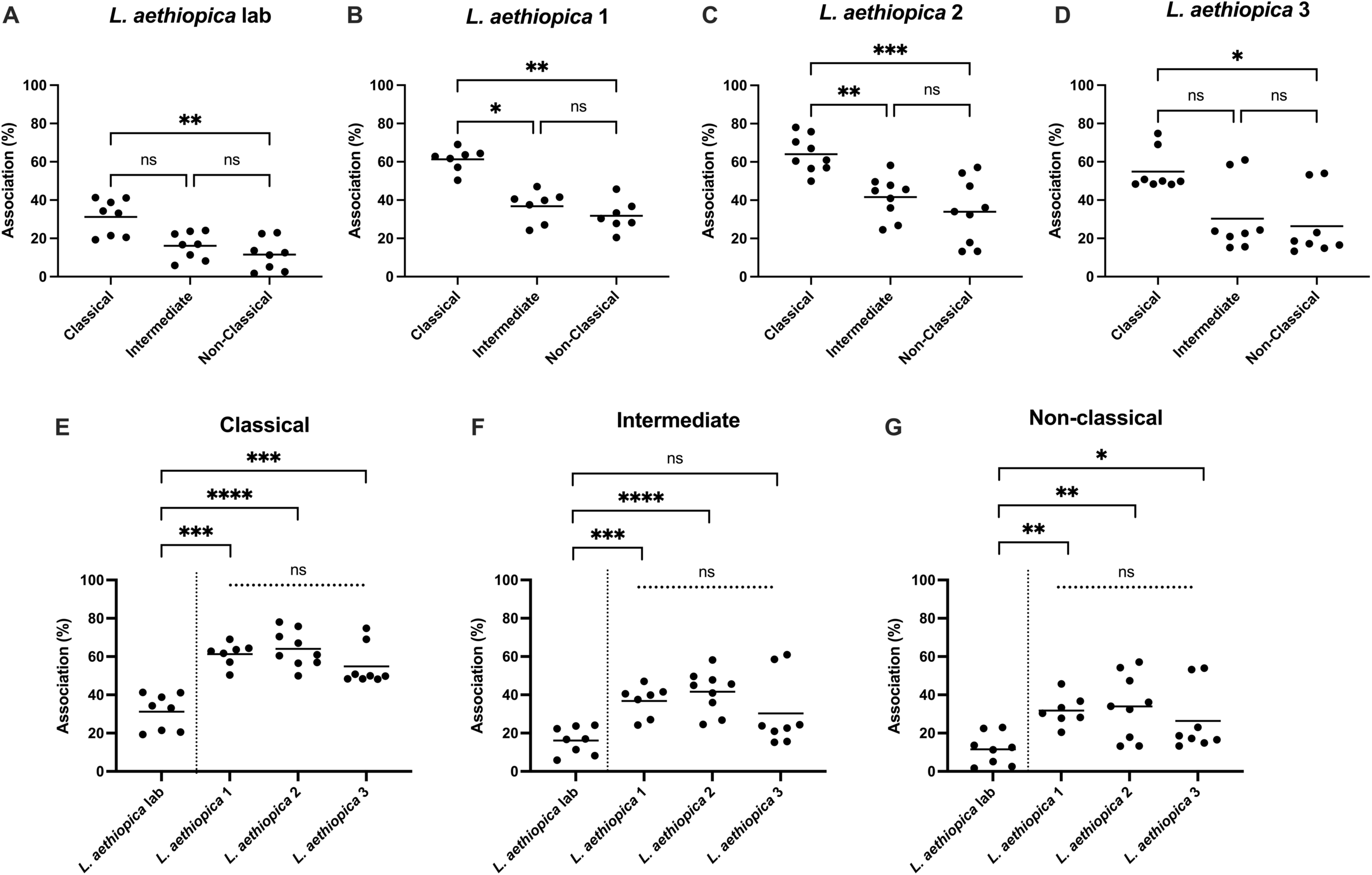
Association of monocyte subsets with *L. aethiopica*. Primary human monocytes were co-incubated for 2 hours with *L. aethiopica^FR^* at an MOI of 1:10. The association of each monocyte subset with parasites was determined by flow cytometry. For each *L. aethiopica* isolate, association was compared between the three monocyte subsets (**A**-**D**), and for each monocyte subset, association with the different *L. aethiopica* isolates was compared (**E**-**G**). *L. aethiopica* lab: n=8, *L. aethiopica* 1: n=7, *L. aethiopica* 2: n=9, *L. aethiopica* 3: n=8. Statistical differences between monocyte subsets were determined using a Kruskal-Wallis test (A-D), between the clinical isolates using a Kruskal-Wallis test (**E-G**), and between *L. aethiopica* lab and each clinical isolate using a Mann-Whitney test (**E-G**). Data summaries and statistical analyses are shown in Tables S2-S4.

We also compared the association between each monocyte subset and the different clinical *L. aethiopica* isolates and show that there was no significant difference in association (Figure 1E-G and Table S3). However, the percentages of monocytes associated with *L. aethiopica* lab were significantly lower than with the clinical isolates, except for the intermediate monocytes co-incubated with *L. aethiopica* 3 (*p*=0.1049) (Figure 1E-G and Table S4).

The flow cytometry experiments described above showed that primary human monocytes have the ability to associate with all *L. aethiopica* parasites. However, the co-localisation of flow cytometry signals (monocytes and *L. aethiopica*) does not necessarily mean internalisation of the parasite. Therefore, confocal microscopy was used to demonstrate that monocytes have the ability to phagocytose *L. aethiopica*. Results presented in Figure 2A-E and Figure S2 confirm that *L. aethiopica* are internalised by monocytes.

**Figure 2-.**
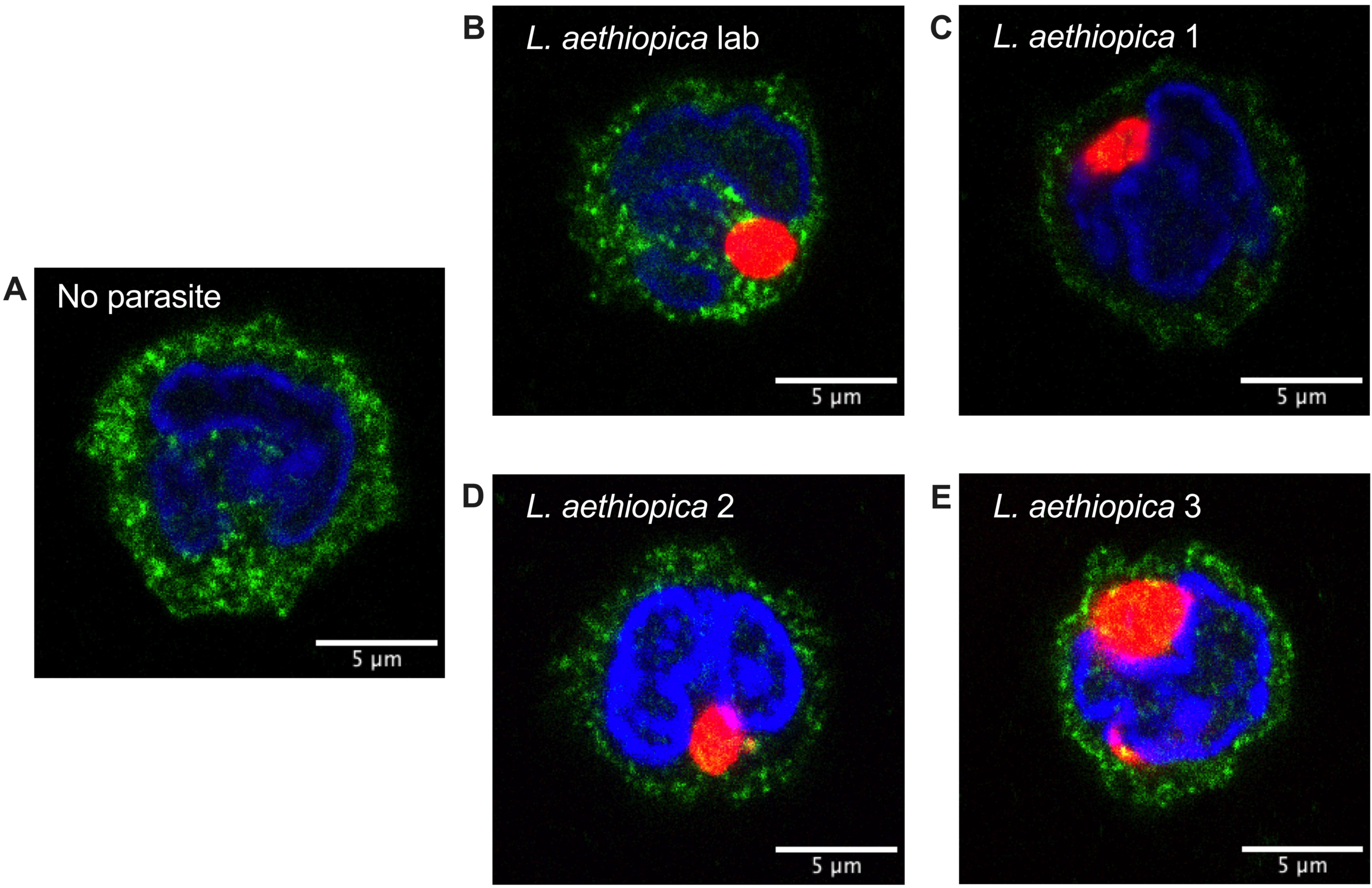
Phagocytosis of *L. aethiopica* by monocytes. Primary human monocytes were co-incubated with *L. aethiopica^FR^* for 2 hours at an MOI of 1:10. Following staining with primary and secondary antibodies, monocytes were visualised using confocal microscopy. Images are shown for monocytes incubated without parasites (**A**), and for monocytes incubated with *L. aethiopica* lab (**B***), L. aethiopica* 1 (**C**), *L. aethiopica* 2 (**D**) and *L. aethiopica* 3 (**E**). One representative image is shown for each parasite, out of three experimental repeats.

### ROS production

Next, we investigated whether the different monocyte subsets produced an oxidative burst in response to co-incubation with the parasites. Results presented in Table S5 show that the mean MFIs of ROS in the unassociated and the associated monocyte subsets were always higher as compared to baseline, however it was not always significant. It was also higher in the associated monocytes than in the unassociated monocytes, although it was not significant for the classical subset incubated with *L. aethiopica* 1 (*p*=0.0584, Table S5).

All three different monocyte subsets associated with *L. aethiopica* produced similar levels of ROS (Figure 3A-D, Table S6). Of note, unassociated classical monocytes produced higher levels of ROS than unassociated non-classical monocytes with *L. aethiopica* lab (Table S6). For each monocyte subset, there was no difference in ROS production in response to all parasite clinical isolates (Table S7). When compared to *L. aethiopica* lab, unassociated classical and non-classical monocytes produced significantly higher levels of ROS following co-incubation with *L. aethiopica* 3 (Table S8).

**Figure 3-.**
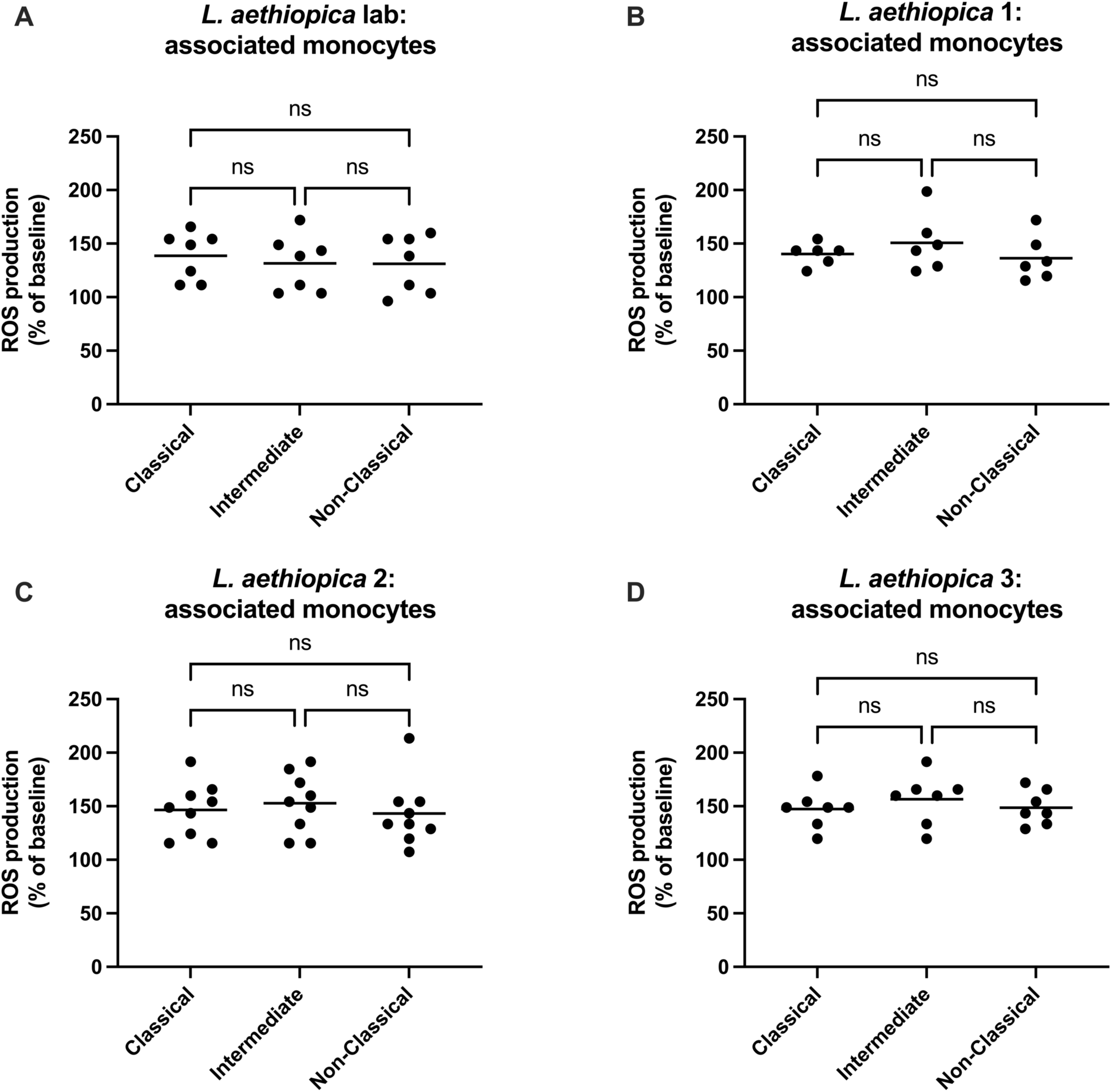
Comparing ROS production by different monocyte subsets. Primary human monocytes were co-incubated with *L. aethiopica^FR^* for 2 hours at an MOI of 1:10. ROS production by monocytes was measured by flow cytometry using a ROS-ID^TM^ Total ROS detection kit. ROS production (as a percentage of baseline ROS production) was compared between the three monocyte subsets for *L. aethiopica* lab (**A***), L. aethiopica* 1 (**B**), *L. aethiopica* 2 (**C**) and *L. aethiopica* 3 (**D**). *L. aethiopica* lab: n=8, *L. aethiopica* 1: n=6, *L. aethiopica* 2: n=9, *L. aethiopica* 3: n=8. Statistical differences between monocyte subsets were determined using a Kruskal-Wallis test. Data summaries and statistical analyses are shown in Table S6.

### Chemokine and cytokine production

#### Chemokines

The levels of eotaxin, eotaxin-3, IP-10, MDC and TARC were below the detection limit. In contrast, IL-8, MCP-1, MCP4, MIP-1α and MIP-1β were significantly upregulated in response to all parasite isolates (Table S9). No significant differences were observed in the production of chemokines following co-incubation with the three clinical isolates (Figure 4 and Table S10). Notably, the levels of MIP-1α and MIP-1β were significantly lower in the supernatants of monocytes co-incubated with *L. aethiopica* lab as compared to *L. aethiopica* 2 and 3 (Figure 4 and Table S11).

**Figure 4-.**
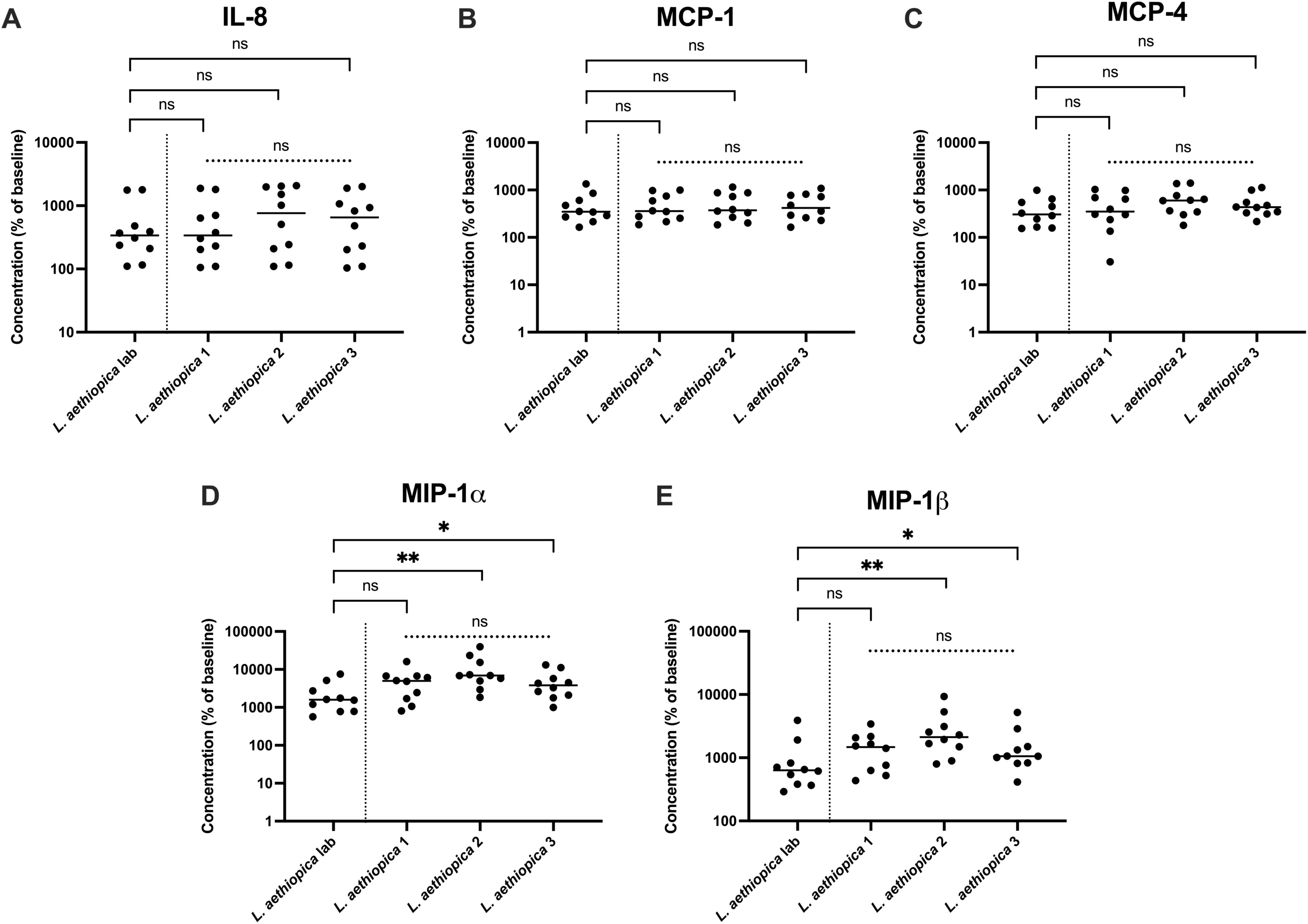
Chemokine production in response to co-incubation with the four *L. aethiopica* isolates. Primary human monocytes were co-incubated with *L. aethiopica* at an MOI of 1:10. After 2 hours, supernatants from each tube were collected. Supernatants were analysed with a V-PLEX Chemokine Panel 1 Human Kit (Meso Scale Diagnostics) to determine the concentration of 10 different chemokines. The concentrations of five chemokines were compared in response to the different parasite isolates: IL-8 (**A**), MCP-1 (**B**), MCP-4 (**C**), MIP-1α (**D**), and MIP-1β (**E**). n=10 for all parasite isolates. Statistical differences between *L. aethiopica* lab and each clinical isolate were determined using a Mann-Whitney test, and between the three clinical isolates using a Kruskal-Wallis test. Data summaries and statistical analyses are shown in Tables S10 and S11.

#### Cytokines

IFN-γ, IL-2, IL-4, IL-12p70, and IL-13 were undetectable. By contrast, the levels of IL-1β, IL-6, IL-10 and TNF-α were significantly increased in response to all parasite isolates, except IL-6 with *L. aethiopica 1* (*p*=0.0982, Table S12). There were no significant differences in cytokine production between the three clinical isolates (Figure 5 and Table S13). However, the levels of IL-1β and TNF-α were significantly lower following co-incubation with *L. aethiopica* lab compared to *L. aethiopica* 2 (Figure 5 and Table S14).

**Figure 5-.**
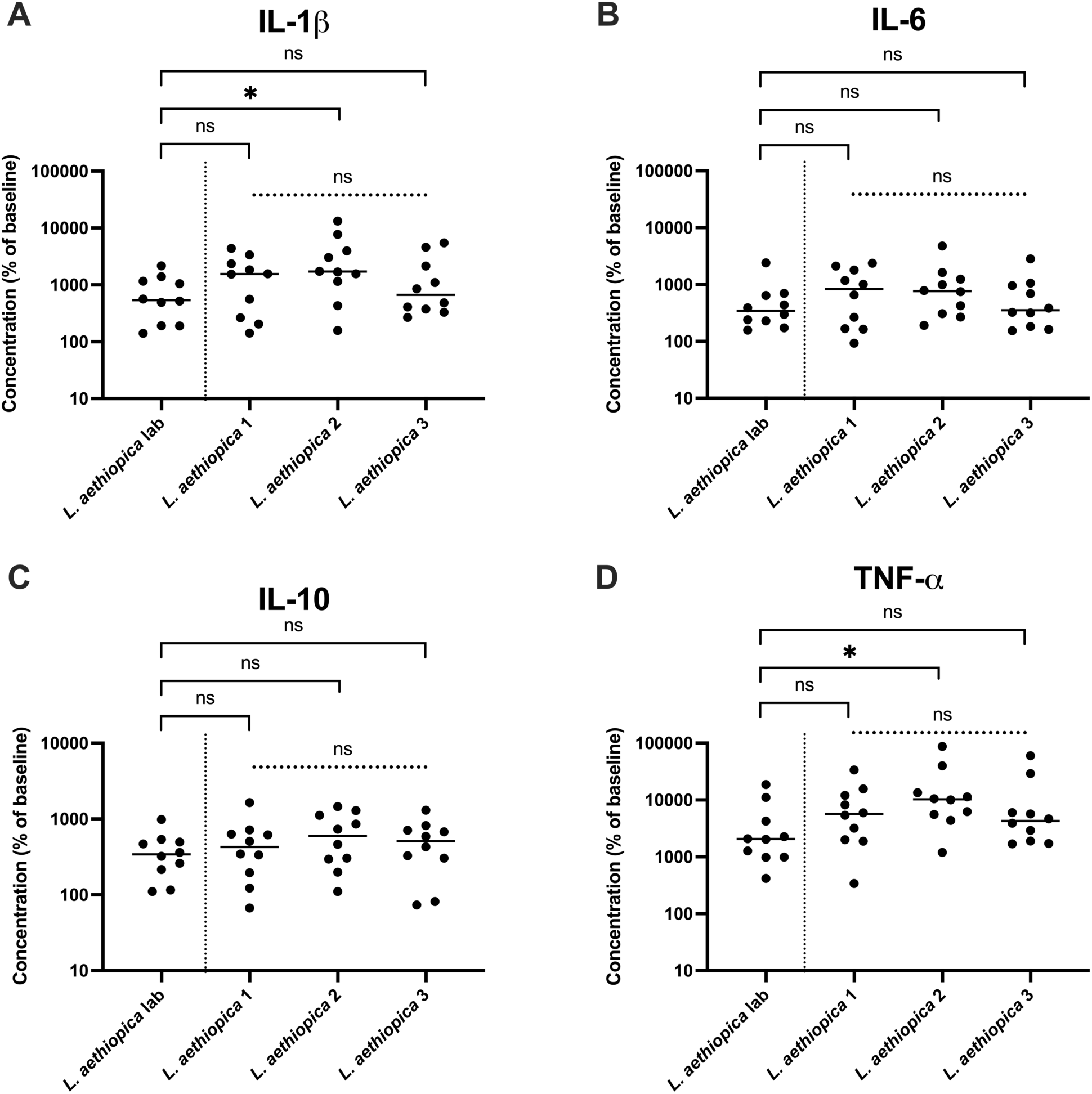
Cytokine production in response to co-incubation with the four *L. aethiopica* isolates. Primary human monocytes were co-incubated with *L. aethiopica* at an MOI of 1:10. After 2 hours, supernatants from each tube were collected. Supernatants were analysed with a V-PLEX Proinflammatory Panel 1 Human Kit (Meso Scale Diagnostics) to determine the concentration of 10 different cytokines. The concentrations of four cytokines were compared in response to the different parasite isolates: IL-1β (**A**), IL-6 (**B**), IL-10 (**C**), TNF-α (**D**). n=10 for all parasite isolates. Statistical differences between *L. aethiopica* lab and each clinical isolate were determined using a Mann-Whitney test, and between the three clinical isolates using a Kruskal-Wallis test. Data summaries and statistical analyses are shown in Tables S13 and S14.

## DISCUSSION

When feeding, sand flies lacerate the skin of the host to form a pool of blood, into which *Leishmania* parasites are deposited [18, 40]. Monocytes are therefore likely to be one of the first cells to encounter parasites. However, little is known about the interactions of monocytes with *L. aethiopica*. Unlike many other CL-causing *Leishmania* species, there are no established animal models for *L. aethiopica* infection [41, 42]. Therefore, here we set up an *in vitro* model, using primary human monocytes. We measured phagocytosis, ROS production and release of chemokines and cytokines in response to co-incubation with *L. aethiopica* and compared those responses to three different recently isolated *L. aethiopica* and a long-term cultured laboratory isolate.

Our results show that all four parasite isolates associated with primary human monocytes. However, some monocytes remained unassociated; this might be due to the short incubation time. Interestingly, while there was no difference between the clinical isolates, association with each of the three clinical isolates was significantly higher than with *L. aethiopica* lab. Phagocytosis of *L. aethiopica* parasites and their entry into monocytes was confirmed using confocal microscopy.

The ability of primary human monocytes to associate with *Leishmania* parasites has already been shown, using different approaches, and using different incubation times as well as different multiplicities of infection (MOIs). This includes whole blood [43, 44], PBMCs [27, 45, 46] and purified human peripheral blood monocytes [32, 47]. Some of the studies specify the parasite stage, e.g. metacyclic parasites; however, an approach to enrich for metacyclic promastigotes is not described [43, 44, 46, 47]. To ensure consistency in the parasite stages that we used, promastigotes were grown to stationary phase and PNA was used in order enrich for metacyclic parasites. Additionally, *L. aethiopica* parasites were kept in culture for a maximum of three weeks. The time parasites are kept in culture is of crucial importance. Indeed, several studies have shown that after *in vitro* culture, parasites lose their virulence as shown by decreased parasites loads *in vivo* and *in vitro* [48–50]. Altered gene and protein expression profiles, as well as chemokine and cytokine production, were reported following extensive *in vitro* passages [51]. In particular, glycoprotein (*GP)63* and *lipophosphoglycan* (*LPG)2* expression were shown to be reduced [52, 53]. Both GP63 and LPG are two of the main surface molecules on *Leishmania* parasites that play a crucial role in phagocytosis [54–56]. Therefore, it is likely that prolonged culture of *L. aethiopica* lab may have resulted in changes in LPG and GP63, leading to the observed differences in association, compared to the clinical *L. aethiopica* isolates.

A recent study assessed the ability of immune cells to associate with GFP-transfected *L. aethiopica* lab [57]. The authors used an MOI of 10:1 (parasites to leukocytes) and reported that an estimated 20% of associated cells were CD14+ cells, as gated in the whole population of GFP+ cells in whole blood, at both 4- and 24-hour timepoints. Since the percentage of monocytes varies greatly between individuals [58], this method does not allow for consistency in MOI between experiments. However, no study has previously shown the association of purified monocytes with *L. aethiopica*.

Our results show that monocytes associated with *L. aethiopica* promastigotes upregulated ROS, as compared to monocytes cultured in the absence of parasites. Incubation with the three clinical isolates, but not *L. aethiopica* lab, also resulted in increased ROS in unassociated monocytes as compared to baseline. ROS MFI in associated monocytes was higher than in unassociated monocytes with all four parasite isolates.

Human monocytes have been shown to produce ROS in response to multiple *Leishmania* species [28, 31, 32, 47, 59]. However, the set-up of these experiments varied greatly, from the incubation time, the parasite species, their life cycle stages, to the ways ROS was measured. Some of the studies have shown that ROS production contributes to killing of the parasite [28, 31, 32]. Yet, others have identified defence mechanisms used by *Leishmania* to inhibit ROS production by phagocytes [60–62]. Of note, our results showed that unassociated monocytes produced an oxidative burst with the clinical isolates of *L. aethiopica*, but not with *L. aethiopica* lab. This could be caused by differences in the surface molecules of *L. aethiopica* lab, especially in LPG and GP63, as discussed above.

Our results showed that five different chemokines were detectable in the supernatants of monocytes incubated with *L. aethiopica* – MCP-1, MCP-4, MIP-1α, MIP-1β, and IL-8. There were higher levels of MIP-1α and MIP-1β following incubation with the clinical isolates than *L. aethiopica lab*. The lower levels of these chemokines in response to *L. aethiopica* lab could result from altered cell surface molecules due to the prolonged time of the parasite isolate in culture.

Little is known about the role of MCP-1 and MCP-4 in leishmaniasis. Several *in vitro* studies using murine and human monocytes and macrophages have shown that the addition of MCP-1 results in increased parasite killing [31, 63, 64]. The *in vitro* production of MIP-1α and MIP-1β by monocytes in response to *Leishmania* parasites has not been previously demonstrated. MIP-1α and MIP-1β were also elevated in the supernatants of PBMCs incubated with *L. amazonensis* in both mouse and human macrophages [63, 64]. In addition to activating the monocytes to kill the parasites, MCP-1, MCP-4, MIP-1α, MIP-1β and IL-8 may contribute to the recruitment of more immune cells and therefore impact on disease development.

Our results also show that production of TNF-α, IL-1β, IL-6 and IL-10 were increased in response to *L. aethiopica*. In our study, IL-6 and IL-10 were similar between all four *L. aethiopica* isolates. Little is known about the production of both cytokines by monocytes co-incubated with *Leishmania* parasites. Two studies have documented the production of IL-6 in response to *L. aethiopica*; however, it was by total PBMCs in response to soluble *Leishmania* antigen (SLA) [65, 66]. Two studies have shown the production of IL-10 by monocytes in response to *L. braziliensis* [46, 67]. TNF-α and IL-1β were increased in response to *L. aethiopica* 2, compared to *L. aethiopica* lab. The production of TNF-α by monocytes in response to pathogens has been extensively documented [68–70], but not to *L. aethiopica*. There are conflicting reports on the production of TNF-α by monocytes in response to *Leishmania* parasites, showing parasite killing or contribution to pathology [71, 72]. Specifically in CL caused by *L. aethiopica*, TNF-α has been shown to be produced by the PBMCs of CL patients in response to incubation with *L. aethiopica* SLA, although the cellular source was not identified [73, 74]. Little is known about the role of IL-1β produced by monocytes in response to *Leishmania* parasites. Santos *et al.* identified intermediate monocytes as the main producers of IL-1β. The authors showed that IL-1β did not contribute to *L. braziliensis* killing by human monocyte-derived macrophages, and suggested that instead, the IL-1β produced contributed to the pathology of CL [75].

It has to be noted that the purity of the monocyte population used in our study was >80% and we can therefore not exclude that other cells contributed to the levels of chemokines and cytokines detected.

Eotaxin, eotaxin-3, IP-10, MDC, TARC, IFN-γ, IL-2, IL-4, IL-8, IL-12p70, and IL-13 were not detectable in the supernatants of activated monocytes. It is possible that these chemokines and cytokines were still produced but were captured by cell surface receptors, or that they were produced at levels below the detection limit.

Although there were no significant differences between clinical isolates in the effector functions measured in this study, there were some differences between *L. aethiopica* lab and some of the clinical *L. aethiopica*; such as the production of TNF-α between *L. aethiopica* lab and *L. aethiopica* 2, but not *L. aethiopica* 1 and *L. aethiopica* 3; this suggests that there may still be differences between the clinical isolates. And indeed, while the three clinical *L. aethiopica* were isolated from LCL patients who had similar lesions, these patients were of different ages, at different times after the onset of symptomatic disease, and had different numbers and sizes of lesions.

One of the main limitations was that we did not measure survival or killing of the parasites. This is usually done by counting intracellular parasites inside the monocytes. However, it is not possible to assess whether these parasites will survive or be killed.

Another method that is commonly used is the transformation assay, where parasites are washed away after co-incubation with the monocytes and the released intracellular parasites are allowed to grow in culture and counted [76]. However, in our experience it has proven impossible to completely eliminate extracellular parasites after the 2-hour incubation, whether by extensive washing or by disrupting the parasite cell membrane using sodium dodecyl sulfate.

In summary, our study describes an *in vitro* model to investigate monocyte effector functions in response to *L. aethiopica*. Our results emphasise that care should be taken when using parasites that have been passaged *in vitro* for a long period of time.

This model could be applied to *L. aethiopica* isolates from patients with LCL, DCL or MCL and might contribute to elucidating why *L. aethiopica* causes different clinical presentations.

## ACKNOWLEDGEMENTS

The authors are thankful to the staff of Nefas Mewcha Hospital for their enthusiastic collaboration during the data collection of this study.

ECC is funded by a Wellcome Trust Studentship (PS3750). This research is jointly funded by the UK Medical Research Council (MRC) and the Foreign Commonwealth and Development Office (FCDO) under the MRC/FCDO Concordat agreement (MR/R021600/1) (EY, JAC, PK). JAC is funded by Wellcome via core funding of the Wellcome Sanger Institute (grant 206194).

---

**Figure S1-.**
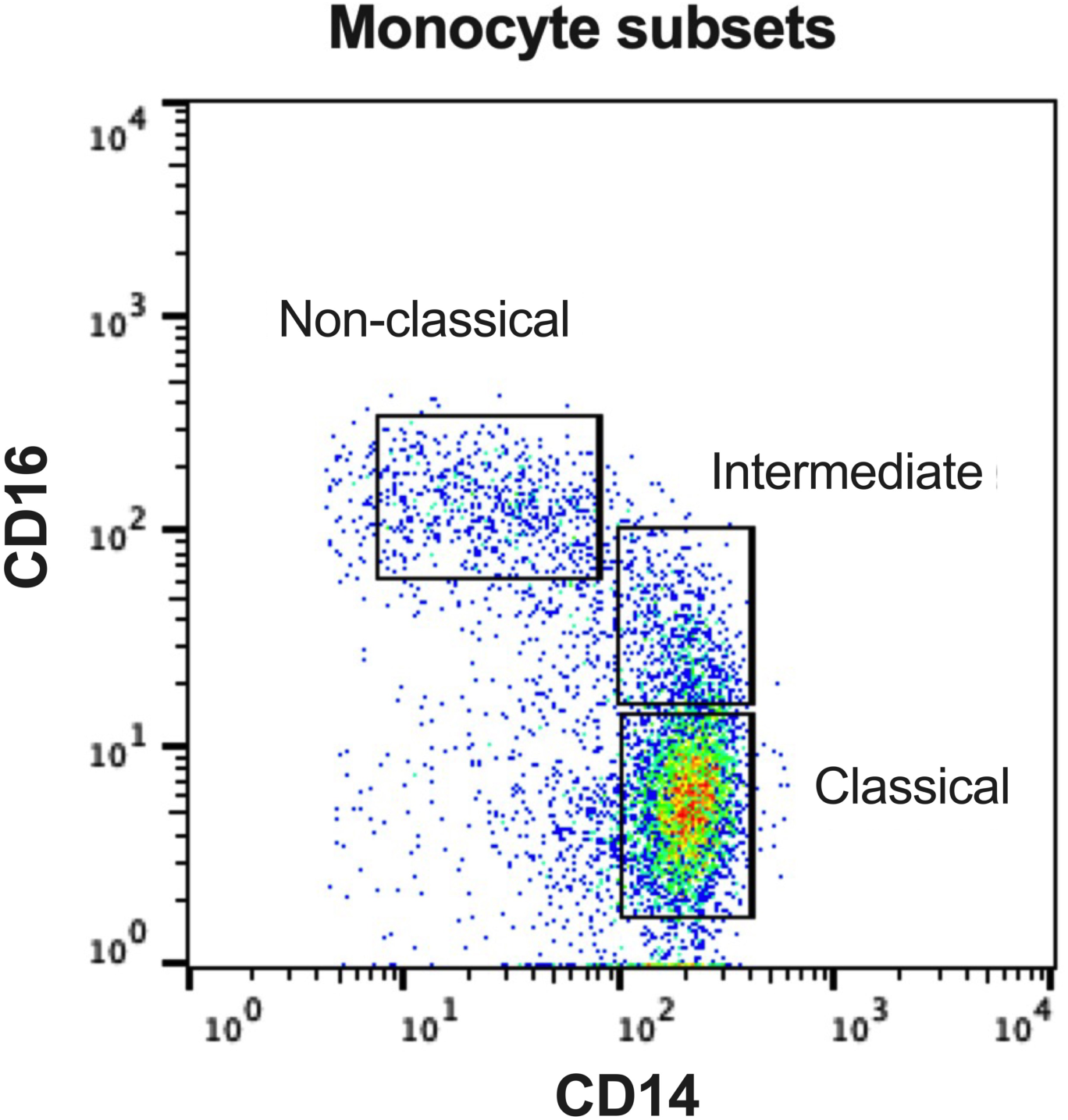
Gating for monocyte subsets. Monocytes were stained with anti-CD14^PE^ and anti-CD16^eFluor™^ ^450^ and divided into subsets according to the following definitions: classical monocytes as CD14^++^ CD16^−^, intermediate monocytes as CD14^++^ CD16^+^ and non-classical monocytes as CD14^+^ CD16^+^.

**Figure S2-.**
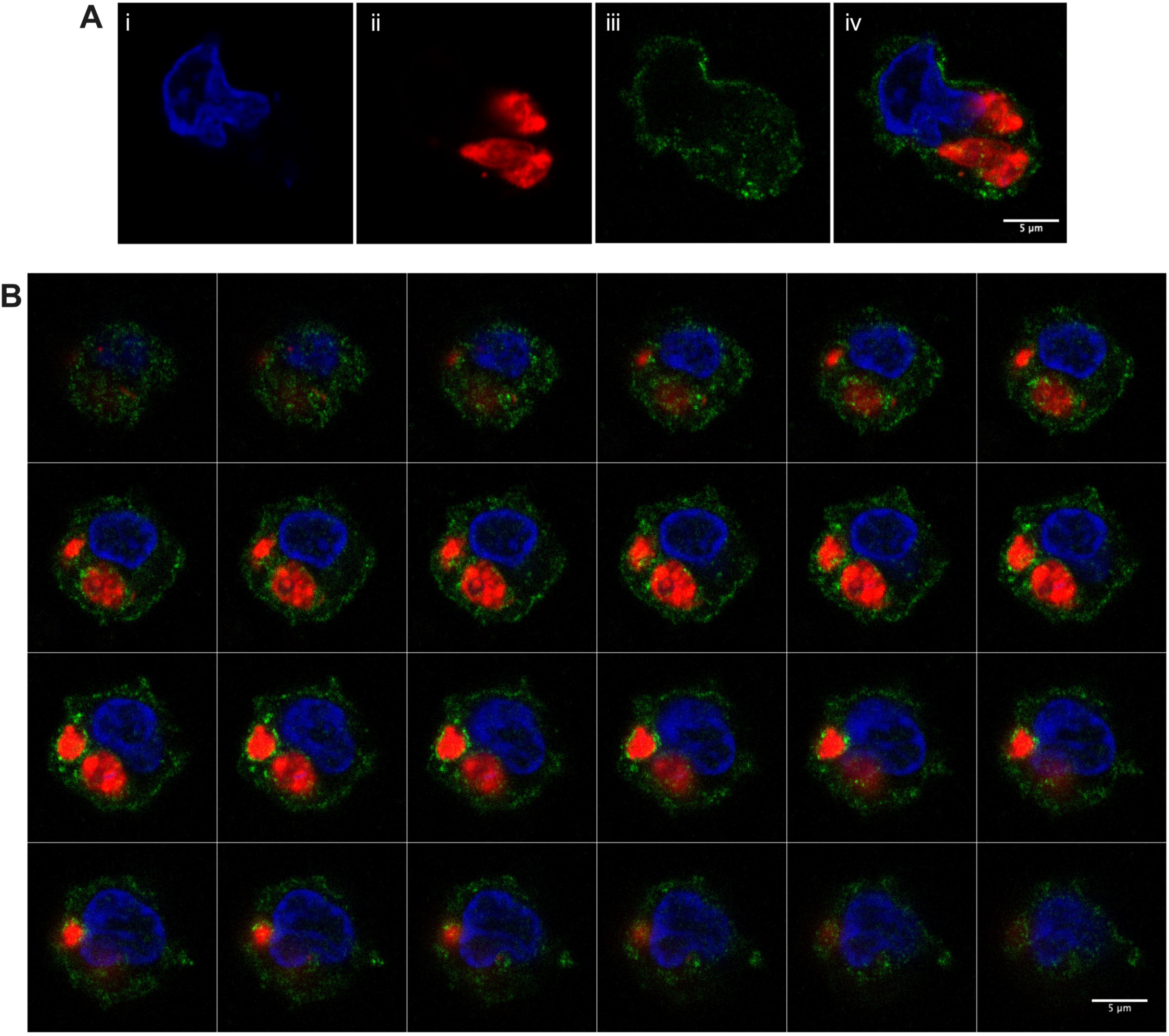
Phagocytosis of *L. aethiopica* promastigotes by primary human monocytes. Primary human monocytes were co-incubated for 2 hours with *L. aethiopica^FR^*at an MOI of 1:10. Following the incubation, monocytes were washed and transferred onto a microscope slide, where they were fixed and stained using anti-CD14 and DAPI. The images show staining for three markers: monocyte surface (anti-CD14^FITC^; green), nuclei (DAPI; blue) and *L. aethiopica* (Far Red dye; red). **A.** The three different channels separately (**i-iii**) and combined (**iv**), for a monocyte infected with *L. aethiopica* 1. **B.** A representative series of 24 z-stack images for a monocyte infected with *L. aethiopica* 1.

**Table S1:** PNA-recovery in stationary phase population: comparison of *L. aethiopica* lab with each clinical isolate.

| *L. aethiopica* | PNA- recovery | PNA- recovery with *L. aethiopica* lab | *p* value* |
| --- | --- | --- | --- |
| 1 | 8.64±2.50 | 3.48±0.72 | **0.0079** |
| 2 | 8.82±1.11 |  | **0.0079** |
| 3 | 10.30±1.52 |  | **0.0079** |

Stationary phase *L. aethiopica* promastigotes were enriched for metacyclic parasites using PNA agglutination as described in Materials and Methods. PNA- recovery (%) is shown as shown as mean ± standard deviation (SD). n=5 for all parasite isolates. Statistical differences between *L. aethiopica* lab and each of the clinical isolates were determined using a Mann-Whitney test\*.

**Table S2:** Comparing association between monocyte subsets, for each *L. aethiopica* isolate.

| *L. aethiopica* | Monocyte subset | Association | *p* value** |  | *p* value*** |
| --- | --- | --- | --- | --- | --- |
| lab | Classical | 31.2±9.4 | **0.0070** | C vs I | 0.1017 |
|  | Intermediate | 16.2±7.1 |  | C vs NC | **0.0063** |
|  | Non-classical | 11.5±8.2 |  | I vs NC | >0.9999 |
| 1 | Classical | 61.3±6.0 | **<0.0001** | C vs I | **0.0153** |
|  | Intermediate | 36.8±8.2 |  | C vs NC | **0.0012** |
|  | Non-classical | 31.8±7.9 |  | I vs NC | >0.9999 |
| 2 | Classical | 64.0±9.4 | **0.0006** | C vs I | **0.0089** |
|  | Intermediate | 41.6±10.9 |  | C vs NC | **0.0009** |
|  | Non-classical | 34.0±16.8 |  | I vs NC | >0.9999 |
| 3 | Classical | 54.9±10.6 | **0.0388** | C vs I | 0.2313 |
|  | Intermediate | 30.3±18.5 |  | C vs NC | **0.0400** |
|  | Non-classical | 26.3±17.1 |  | I vs NC | >0.9999 |

Primary human monocytes were co-incubated for 2 hours with *L. aethiopica*<sup>FR</sup> promastigotes at an MOI of 1:10. The association (%) of each subset with parasites was determined by flow cytometry, and subsets were compared for each parasite isolate. *L. aethiopica* lab: n=8, *L. aethiopica* 1: n=7, *L. aethiopica* 2: n=9, and *L. aethiopica* 3: n=8. Statistical differences were determined using a Kruskal-Wallis test\*\* and Dunn's multiple comparisons test\*\*\*.

**Table S3:** Comparing association with the three clinical isolates of *L. aethiopica*, for each of the monocyte subsets.

| Monocyte subset | *L. aethiopica* | Association | *p* value** |
| --- | --- | --- | --- |
| Classical | 1 | 61.3±6.0 | 0.0986 |
|  | 2 | 64.0±9.4 |  |
|  | 3 | 54.9±10.6 |  |
| Intermediate | 1 | 36.8±8.2 | 0.1209 |
|  | 2 | 41.6±10.9 |  |
|  | 3 | 30.3±18.5 |  |
| Non-classical | 1 | 31.8±7.9 | 0.4202 |
|  | 2 | 34.0±16.8 |  |
|  | 3 | 26.3±17.1 |  |

Primary human monocytes were co-incubated for 2 hours with *L. aethiopica*<sup>FR</sup> promastigotes at an MOI of 1:10. Association (%) was determined by flow cytometry and is shown as mean ± SD. *L. aethiopica* 1: n=7, *L. aethiopica* 2: n=9, and *L. aethiopica* 3: n=8. Statistical differences between the three clinical isolates were determined using a Kruskal-Wallis test\*\*.

**Table S4-.**
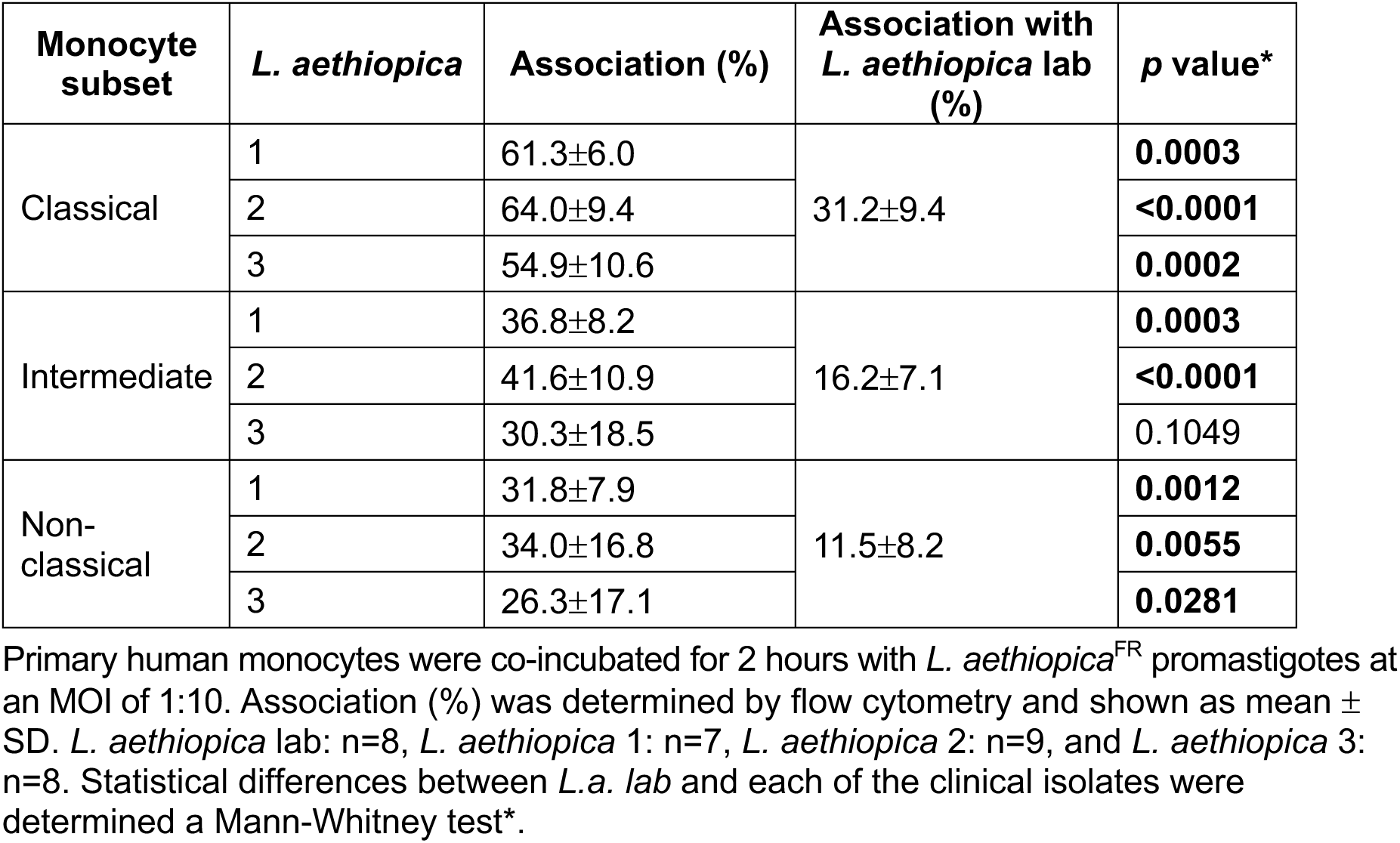
Comparing monocyte association with *L. aethiopica lab* and three clinical isolates of *L. aethiopica*.

| Monocyte subset | *L. aethiopica* | Association (%) | Association with *L. aethiopica* lab (%) | *p* value* |
| --- | --- | --- | --- | --- |
| Classical | 1 | 61.3±6.0 | 31.2±9.4 | **0.0003** |
|  | 2 | 64.0±9.4 |  | **<0.0001** |
|  | 3 | 54.9±10.6 |  | **0.0002** |
| Intermediate | 1 | 36.8±8.2 | 16.2±7.1 | **0.0003** |
|  | 2 | 41.6±10.9 |  | **<0.0001** |
|  | 3 | 30.3±18.5 |  | 0.1049 |
| Non-classical | 1 | 31.8±7.9 | 11.5±8.2 | **0.0012** |
|  | 2 | 34.0±16.8 |  | **0.0055** |
|  | 3 | 26.3±17.1 |  | **0.0281** |

Primary human monocytes were co-incubated for 2 hours with *L. aethiopica*<sup>FR</sup> promastigotes at an MOI of 1:10. Association (%) was determined by flow cytometry and shown as mean ± SD. *L. aethiopica* lab: n=8, *L. aethiopica* 1: n=7, *L. aethiopica* 2: n=9, and *L. aethiopica* 3: n=8. Statistical differences between *L.a. lab* and each of the clinical isolates were determined a Mann-Whitney test\*.

**Table S5:**
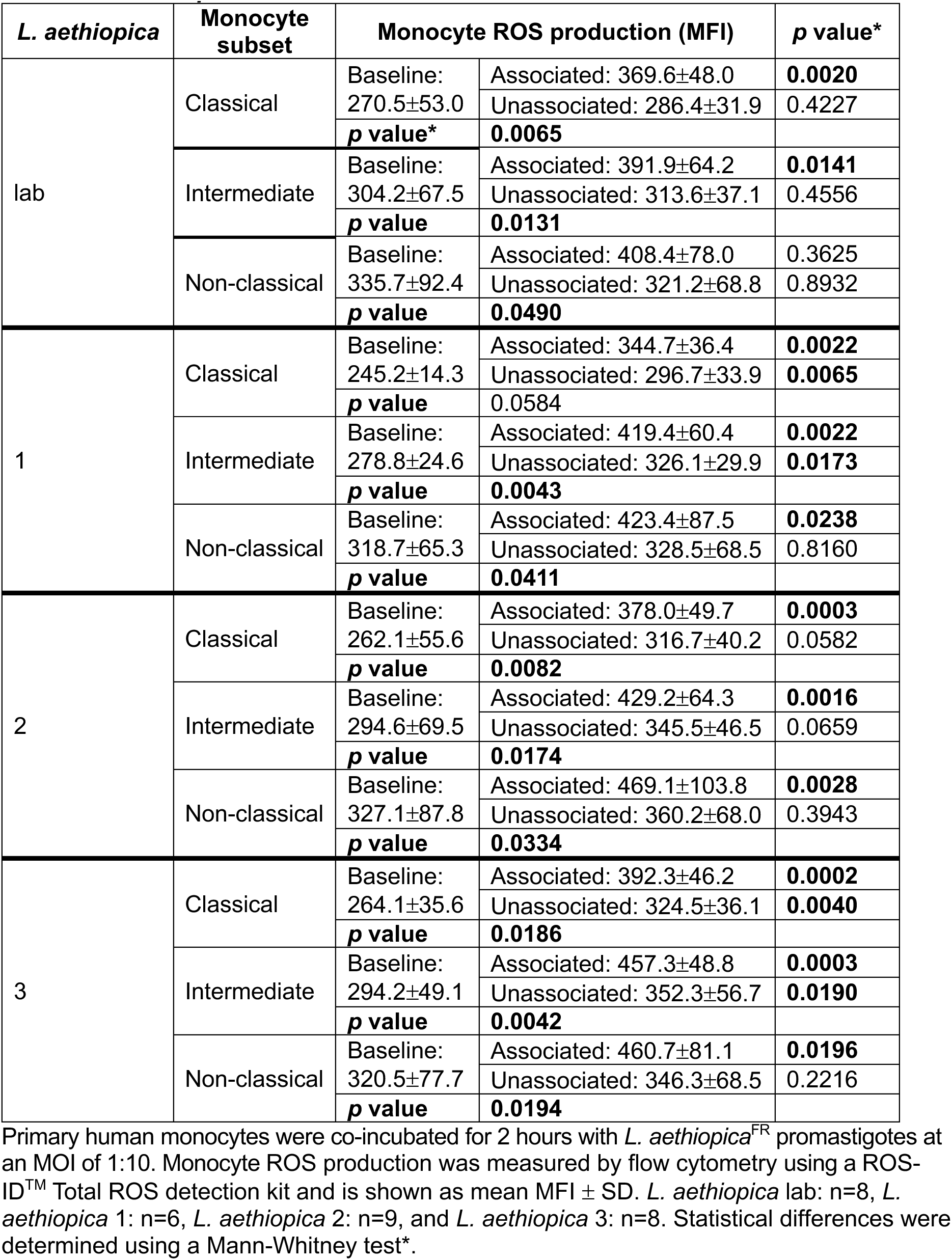
ROS production by monocyte subsets following co-incubation with the different *L. aethiopica* isolates.

**Table S6:** Comparing ROS production by monocyte subsets in response to *L. aethiopica*.

| *L. aethiopica* | Association | Subsets | ROS | *p* value** | Groups | *p* value*** |
| --- | --- | --- | --- | --- | --- | --- |
| lab | Associated | Classical | 138.6±22.4 | 0.3045 | NA |  |
|  |  | Intermediate | 131.6±26.1 |  |  |  |
|  |  | Non-classical | 131.2±26.8 |  |  |  |
|  | Unassociated | Classical | 110.4±17.8 | **0.0187** | C v I | 0.7618 |
|  |  | Intermediate | 107.4±19.5 |  | C v NC | **0.0215** |
|  |  | Non-classical | 98.2±12.8 |  | I v NC | >3646 |
| 1 | Associated | Classical | 140.4±10.3 | 0.5063 | NA |  |
|  |  | Intermediate | 150.6±26.9 |  |  |  |
|  |  | Non-classical | 136.4±20.9 |  |  |  |
|  | Unassociated | Classical | 120.8±8.3 | 0.5063 | NA |  |
|  |  | Intermediate | 115.1±8.6 |  |  |  |
|  |  | Non-classical | 105.8±8.2 |  |  |  |
| 2 | Associated | Classical | 146.6±25.2 | 0.6517 | NA |  |
|  |  | Intermediate | 152.9±27.6 |  |  |  |
|  |  | Non-classical | 143.2±30.4 |  |  |  |
|  | Unassociated | Classical | 122.8±17.3 | 0.0810 | NA |  |
|  |  | Intermediate | 125.3±20.2 |  |  |  |
|  |  | Non-classical | 115.5±18.8 |  |  |  |
| 3 | Associated | Classical | 147.4±18.1 | 0.5691 | NA |  |
|  |  | Intermediate | 156.6±23.5 |  |  |  |
|  |  | Non-classical | 148.7±16.0 |  |  |  |
|  | Unassociated | Classical | 123.5±10.7 | 0.6838 | NA |  |
|  |  | Intermediate | 121.2±9.5 |  |  |  |
|  |  | Non-classical | 112.7±8.0 |  |  |  |

Primary human monocytes were co-incubated for 2 hours with *L. aethiopica*<sup>FR</sup> promastigotes at an MOI of 1:10. Monocyte ROS production was measured by flow cytometry using a ROS-ID™ Total ROS detection kit, was calculated as % of baseline ROS production, and is shown as mean ± SD. *L. aethiopica* lab: n=8, *L. aethiopica* 1: n=6, *L. aethiopica* 2: n=9, and *L. aethiopica* 3: n=8. Statistical differences between monocyte subsets were determined using a Kruskal-Wallis test\*\*.

**Table S7:** Comparing ROS production by monocyte subsets co-incubated with the clinical isolates.

| Monocyte subset | Association | *L. aethiopica* | ROS | p value** |
| --- | --- | --- | --- | --- |
| Classical | Associated | 1 | 140.4±10.3 | 0.4585 |
|  |  | 2 | 147.8±25.4 |  |
|  |  | 3 | 149.7±18.1 |  |
|  | Unassociated | 1 | 120.8±8.3 | 0.9373 |
|  |  | 2 | 123.2±16.9 |  |
|  |  | 3 | 123.5±10.7 |  |
| Intermediate | Associated | 1 | 151.4±25.8 | 0.6040 |
|  |  | 2 | 149.6±26.2 |  |
|  |  | 3 | 157.8±22.4 |  |
|  | Unassociated | 1 | 117.3±10.4 | 0.5280 |
|  |  | 2 | 123.6±18.7 |  |
|  |  | 3 | 120.1±9.2 |  |
| Non-classical | Associated | 1 | 133.6±17.5 | 0.4812 |
|  |  | 2 | 147.1±29.5 |  |
|  |  | 3 | 146.2±16.1 |  |
|  | Unassociated | 1 | 103.4±8.5 | 0.4432 |
|  |  | 2 | 113.2±17.9 |  |
|  |  | 3 | 109.1±6.0 |  |

Primary human monocytes were co-incubated for 2 hours with *L. aethiopica*<sup>FR</sup> promastigotes at an MOI of 1:10. Monocyte ROS production was measured by flow cytometry using a ROS-ID™ Total ROS detection kit, was calculated as % of baseline ROS production, and is shown as mean ± SD. *L. aethiopica* 1: n=6, *L. aethiopica* 2: n=9, and *L. aethiopica* 3: n=8. Statistical differences between the three clinical isolates were determined using a Kruskal-Wallis test\*\*.

**Table S8-.** Comparison of ROS production by monocyte subsets between *L. aethiopica lab* with each clinical isolate.

| Monocyte subset | Association | *L. aethiopica* | ROS | ROS with *L. aethiopica* lab | *p* value* |
| --- | --- | --- | --- | --- | --- |
| Classical | Associated | 1 | 140.4±10.3 | 139.4±22.3 | 0.8032 |
|  |  | 2 | 147.8±25.4 |  | 0.4954 |
|  |  | 3 | 149.7±18.1 |  | 0.5936 |
|  | Unassociated | 1 | 120.8±8.3 | 108.0±15.7 | 0.0982 |
|  |  | 2 | 123.2±16.9 |  | 0.1191 |
|  |  | 3 | 123.5±10.7 |  | **0.0465** |
| Intermediate | Associated | 1 | 151.4±25.8 | 132.1±25.7 | 0.2128 |
|  |  | 2 | 149.6±26.2 |  | 0.2076 |
|  |  | 3 | 157.8±22.4 |  | 0.0690 |
|  | Unassociated | 1 | 117.3±10.4 | 105.7±16.4 | 0.1695 |
|  |  | 2 | 123.6±18.7 |  | 0.0607 |
|  |  | 3 | 120.1±9.2 |  | 0.0716 |
| Non-classical | Associated | 1 | 133.6±17.5 | 125.9±26.6 | 0.6509 |
|  |  | 2 | 147.1±29.5 |  | 0.6096 |
|  |  | 3 | 146.2±16.1 |  | 0.8275 |
|  | Unassociated | 1 | 103.4±8.5 | 97.4±10.4 | 0.3287 |
|  |  | 2 | 113.2±17.9 |  | 0.0778 |
|  |  | 3 | 109.1±6.0 |  | **0.0298** |

Primary human monocytes were co-incubated for 2 hours with *L. aethiopica*<sup>FR</sup> promastigotes at an MOI of 1:10. P Monocyte ROS production was measured by flow cytometry using a ROS-ID™ Total ROS detection kit, was calculated as % of baseline ROS production, and is shown as mean ± SD. *L. aethiopica* lab: n=8, *L. aethiopica* 1: n=6, *L. aethiopica* 2: n=9, and *L. aethiopica* 3: n=8. Statistical differences between *L. aethiopica* lab and each clinical isolate were determined using a Mann-Whitney test\*.

**Table S9-.** Chemokine production in response to co-incubation with the four *L. aethiopica* isolates.

| Chemokine | *L. aethiopica* | Concentration | Baseline concentration | *p* value* |
| --- | --- | --- | --- | --- |
| MCP-1 | lab | 169.8±145.0 | 58.32±63.41 | **0.0185** |
|  | 1 | 172.6±140.3 |  | **0.0288** |
|  | 2 | 177.4±139.5 |  | **0.0115** |
|  | 3 | 175.0±136.5 |  | **0.0115** |
| MCP-4 | lab | 15.59±12.36 | 4.337±2.064 | **0.0002** |
|  | 1 | 19.65±18.02 |  | **0.0068** |
|  | 2 | 24.28±13.77 |  | **<0.0001** |
|  | 3 | 19.38±8.883 |  | **<0.0001** |
| MIP-1α | lab | 352.0±374.6 | 22.34±27.03 | **<0.0001** |
|  | 1 | 1271±1596 |  | **<0.0001** |
|  | 2 | 1692±1419 |  | **<0.0001** |
|  | 3 | 799.0±1044 |  | **<0.0001** |
| MIP-1β | lab | 430.9±412.2 | 81.66±118.6 | **0.0011** |
|  | 1 | 1187±1844 |  | **0.0005** |
|  | 2 | 2134±3655 |  | **<0.0001** |
|  | 3 | 1013±1743 |  | **0.0002** |
| IL-8 | lab | 785.3±500.3 | 371.3±447.8 | **0.0232** |
|  | 1 | 798.8±453.5 |  | **0.0232** |
|  | 2 | 983.0±314.7 |  | **0.0029** |
|  | 3 | 876.4±373.3 |  | **0.0147** |

Primary human monocytes were co-incubated for 2 hours with *L. aethiopica* promastigotes at an MOI of 1:10. Supernatants were collected and analysed on a V-PLEX Chemokine Panel 1 Human Kit to determine the concentration of 10 different chemokines. Chemokine concentration (pg/mL) is shown as mean ± SD. n=10 for all parasite isolates. Statistical differences were determined using a Mann-Whitney test\*.

**Table S10-.** Comparing chemokine production in response to co-incubation with clinical isolates.

| Chemokine | *L. aethiopica* | Concentration | *p* value** |
| --- | --- | --- | --- |
| MCP-1 | 1 | 494.9±310.3 | 0.9683 |
|  | 2 | 535.5±340.5 |  |
|  | 3 | 517.9±310.8 |  |
| MCP-4 | 1 | 474.6±344.2 | 0.5661 |
|  | 2 | 656.1±423.3 |  |
|  | 3 | 521.5±299.6 |  |
| MIP-1α | 1 | 5172±4493 | 0.0964 |
|  | 2 | 11508±11824 |  |
|  | 3 | 4971±4077 |  |
| MIP-1β | 1 | 1459±932.7 | 0.1289 |
|  | 2 | 2941±2593 |  |
|  | 3 | 1610±1426 |  |
| IL-8 | 1 | 635.6±667.5 | 0.6069 |
|  | 2 | 980.8±848.4 |  |
|  | 3 | 784.3±704.2 |  |

Primary human monocytes were co-incubated for 2 hours with *L. aethiopica* promastigotes at an MOI of 1:10. Supernatants were collected and analysed on a V-PLEX Chemokine Panel 1 Human Kit to determine the concentration of 10 different chemokines. Chemokine concentration was calculated as % of baseline production (%) is shown as mean ± SD. n=10 for all parasite isolates. Statistical differences between the three clinical isolates were determined using a Kruskal-Wallis test\*\*.

**Table S11-.** Comparing chemokine production in response to co-incubation with *L.a. lab* and clinical isolates.

| Chemokine | *L. aethiopica* | Concentration | Concentration with *L. aethiopica* lab | *p* value* |
| --- | --- | --- | --- | --- |
| MCP-1 | 1 | 494.9±310.3 | 491.5±364.4 | 0.9118 |
|  | 2 | 535.5±340.5 |  | 0.7394 |
|  | 3 | 517.9±310.8 |  | 0.7959 |
| MCP-4 | 1 | 474.6±344.2 | 397.2±269.4 | 0.7394 |
|  | 2 | 656.1±423.3 |  | 0.0892 |
|  | 3 | 521.5±299.6 |  | 0.2176 |
| MIP-1α | 1 | 5172±4493 | 2364±2248 | 0.1051 |
|  | 2 | 11508±11824 |  | **0.0021** |
|  | 3 | 4971±4077 |  | **0.0433** |
| MIP-1β | 1 | 1459±932.7 | 1018±1110 | 0.1655 |
|  | 2 | 2941±2593 |  | **0.0039** |
|  | 3 | 1610±1426 |  | **0.0433** |
| IL-8 | 1 | 635.6±667.5 | 577.0±642.5 | 0.9705 |
|  | 2 | 980.8±848.4 |  | 0.4359 |
|  | 3 | 784.3±704.2 |  | 0.6305 |

Primary human monocytes were co-incubated for 2 hours with *L. aethiopica* promastigotes at an MOI of 1:10. Supernatants were collected and analysed on a V-PLEX Chemokine Panel 1 Human Kit to determine the concentration of 10 different chemokines. Chemokine concentration was calculated as % of baseline production (%) is shown as mean ± SD. n=10 for all parasite isolates. Statistical differences between *L. aethiopica* lab and each clinical isolate were determined using a Mann-Whitney test\*.

**Table S12-.** Cytokine production in response to co-incubation with the four *L. aethiopica* isolates.

| **Cytokine** | ***L. aethiopica*** | **Concentration** | **Baseline concentration** | **p value\*\*** |
| --- | --- | --- | --- | --- |
| TNF-α | lab | 134.9±98.14 | 6.137±6.628 | **0.0001** |
| | 1 | 443.9±458.2 | | **0.0001** |
| | 2 | 597.5±395.8 | | **<0.0001** |
| | 3 | 299.7±221.2 | | **<0.0001** |
| IL-1β | lab | 6.984±8.142 | 0.9005±0.9754 | **0.0185** |
| | 1 | 20.94±28.01 | | **0.0147** |
| | 2 | 22.22±23.20 | | **0.0005** |
| | 3 | 9.353±10.82 | | **0.0052** |
| IL-6 | lab | 29.10±43.89 | 6.586±10.40 | **0.0433** |
| | 1 | 88.40±137.6 | | 0.0982 |
| | 2 | 64.95±101.3 | | **0.0147** |
| | 3 | 48.22±96.99 | | **0.0433** |
| IL-10 | lab | 0.9682±0.9523 | 0.2390±0.2181 | **0.0147** |
| | 1 | 1.448±1.693 | | **0.0355** |
| | 2 | 1.567±1.343 | | **0.0068** |
| | 3 | 1.209±1.070 | | **0.0185** |

Primary human monocytes were co-incubated for 2 hours with *L. aethiopica* promastigotes at an MOI of 1:10. Primary human monocytes were co-incubated for 2 hours with *L. aethiopica* promastigotes at an MOI of 1:10. Supernatants were collected and analysed on a V-PLEX Proinflammatory Panel 1 Human Kit to determine the concentration of 10 different cytokines. Cytokine concentration (pg/mL) is shown as mean ± SD. n=10 for all parasite isolates. Statistical differences were determined using a Mann-Whitney test\*.

**Table S13-.** Comparing cytokine production in response to co-incubation with clinical isolates.

| Cytokine | *L. aethiopica* | Concentration | *p* value** |
| --- | --- | --- | --- |
| TNF-α | 1 | 8838±9970 | 0.2842 |
| | 2 | 19028±26340 | |
| | 3 | 11782±18872 | |
| IL-1β | 1 | 1625±1435 | 0.4137 |
| | 2 | 3468±4069 | |
| | 3 | 1603±1901 | |
| IL-6 | 1 | 981.0±858.2 | 0.5764 |
| | 2 | 1138±1359 | |
| | 3 | 708.0±817.3 | |
| IL-10 | 1 | 519.8±455.6 | 0.7159 |
| | 2 | 683.1±482.4 | |
| | 3 | 531.6±372.5 | |

Primary human monocytes were co-incubated for 2 hours with *L. aethiopica* promastigotes at an MOI of 1:10. Supernatants were collected and analysed on a V-PLEX Proinflammatory Panel 1 Human Kit to determine the concentration of 10 different cytokines. Cytokine concentration was calculated as % of baseline production (%) is shown as mean ± SD. n=10 for all parasite isolates. Statistical differences between the three clinical isolates were determined using a Kruskal-Wallis test\*\*.

**Table S14-.** Comparing cytokine production in response to co-incubation with *L.a. lab* and clinical isolates.

| Cytokine | *L. aethiopica* | Concentration | Concentration with *L. aethiopica* lab | *p* value* |
| --- | --- | --- | --- | --- |
| TNF-α | 1 | 8838±9970 | 4417±5926 | 0.2176 |
| | 2 | 19028±26340 | | **0.0185** |
| | 3 | 11782±18872 | | 0.1431 |
| IL-1β | 1 | 1625±1435 | 787.6±651.1 | 0.1655 |
| | 2 | 3468±4069 | | **0.0355** |
| | 3 | 1603±1901 | | 0.6305 |
| IL-6 | 1 | 981.0±858.2 | 568.7±675.5 | 0.5288 |
| | 2 | 1138±1359 | | 0.1051 |
| | 3 | 708.0±817.3 | | 0.7959 |
| IL-10 | 1 | 519.8±455.6 | 388.2±259.0 | 0.5288 |
| | 2 | 683.1±482.4 | | 0.2799 |
| | 3 | 531.6±372.5 | | 0.4359 |

Primary human monocytes were co-incubated for 2 hours with *L. aethiopica* promastigotes at an MOI of 1:10. Supernatants were collected and analysed on a V-PLEX Proinflammatory Panel 1 Human Kit to determine the concentration of 10 different cytokines. Cytokine concentration was calculated as % of baseline production (%) is shown as mean ± SD. n=10 for all parasite isolates. Statistical differences between *L. aethiopica* lab and each clinical isolate were determined using a Mann-Whitney test\*.

